# Revisiting synthetic lethality of Gcn5-related N-acetyltransferase (GNAT) family mutations in *Haloferax volcanii*

**DOI:** 10.1101/2025.02.13.638158

**Authors:** Katherine R. Weber, Brianna Novillo, Julie A. Maupin-Furlow

## Abstract

Lysine acetylation is a post-translational modification that occurs in all domains of life, highlighting its evolutionary significance. Previous genome comparison identified three Gcn5-related N-acetyltransferase (GNAT) family members as lysine acetyltransferase homologs (Pat1, Pat2, and Elp3) and two deacetylase homologs (Sir2 and HdaI) in the halophilic archaeon *Haloferax volcanii*, with *elp3* and *pat2* proposed as a synthetic lethal gene pair. Here we advance these findings by performing single and double mutagenesis of *elp3* with the *pat1* and *pat2* lysine acetyltransferase gene homologs. Genome sequencing and PCR screens of these strains reveal successful generation of *Δelp3, Δpat1Δelp3*, and *Δpat2Δelp3* mutant strains. Although these mutant strains exhibited a reduced growth rate compared to the parent, they remained viable. Overall, this study provides genetic evidence that *elp3* and *pat2*, while impacting cell growth, are not a synthetic lethal gene pair as previously reported.

**IMPORTANCE:** Here we reveal by whole genome sequencing that the GNAT family gene homologs *elp3* and *pat2* can be deleted in the same *H. volcanii* strain. Beyond the targeted deletions, minimal differences between the parent and *Δelp3 Δpat2* mutant were observed suggesting that suppressor mutations are not responsible for our ability to generate this double mutant strain. Elp3 and Pat2, thus, may not share as close a functional relationship as implied by earlier study. Our finding is significant as Elp3 is thought to function in acetylation in tRNA modification, while Pat2 likely functions in the lysine acetylation of proteins.

## INTRODUCTION

Post-translational modifications (PTMs) are chemical modifications that occur after protein synthesis, influencing the charge, structure, and function of proteins (1). Among PTMs, lysine acetylation is considered an ancient form that is evolutionarily conserved in all domains of life (2). It regulates key biological processes, including metabolism and chromatin structure, and is often responsive to external stimuli such as nutrient availability and oxidative stress (3-5).

Lysine acetylation involves the transfer of an acetyl group from the metabolic intermediates acetyl-coenzyme A (acetyl-CoA) or acetyl-phosphate (acetyl-P) to the lysine residue of a target protein (6). This reaction can occur non-enzymatically or enzymatically by lysine acetyltransferases, with lysine deacetylases catalyzing the removal of the acetyl group, offering opportunities for regulatory control (3). While research has examined lysine acetylation extensively in the context of histone modification and bacterial metabolism (3-5), the role of this PTM in archaeal cell biology is less studied.

*Haloferax volcanii* (*Hv*) is a hypersaline-adapted archaeon that tolerates harsh environmental conditions such as ultraviolet (UV) irradiation (7), desiccation (8), extreme temperatures and pH (9, 10), heavy metals (11), and strong oxidants (12, 13). Proteomic and genetic studies of *H. volcanii* reveal a correlation between oxidative stress and lysine acetylation (12-14). Previous genome comparison identified three histone lysine acetyltransferase (HAT) homologs of the Gcn5-related N-acetyltransferase (GNAT) family, Pat1, Pat2 and Elp3, and two histone deacetylase (HDAC) homologs including the NAD^+^-dependent class III Sir2 and the zinc-dependent class II Hdal (15). While *Hv*Elp3 is of the GNAT family, it is also related to and annotated as a tRNA carboxymethyluridine synthase (16). In eukaryotic systems, Elp3 is the catalytic subunit of a the Elongator (Elp) complex, which catalyzes uridine modifications at the wobble position (17). This subunit contains two putative domains: radical S-adenosylmethionine (SAM) at the N-terminus and the histone acetyltransferase (HAT or GNAT) domain at the C-terminus (18, 19). A study in the archaea *Methanocaldococcus infernus* aimed to perform an *in vitro* reconstitution of radiolabeled acetyl-CoA and synthetic tRNA (20). The results suggest the C-terminal GNAT domain of Elp3 is utilized for adding a carboxymethyl (cm^5^) group to uridine at the wobble position, reproposing Elp3 is not a lysine acetyltransferase but instead is involved with wobble uridine tRNA modification (20).

Due to the close evolutionary relationship of archaea with eukaryotes, *H. volcanii* has emerged as a model organism widely used in genetic, molecular, and biochemical research to provide valuable insights into cellular mechanisms of survival under extreme environmental conditions (21). Previous efforts aimed to generate a *Δpat2 Δelp3* double mutant suggested that these genes are involved in synthetic lethal interaction, in which their products share the same targets, and deemed essential for *H. volcanii* viability (19). Here we provide genetic and phenotypic evidence in *H. volcanii* that *elp3* can be deleted in combination with either *pat1* or *pat2*, thus, supporting a conclusion that a *Δpat2 Δelp3* double mutation is not lethal.

## MATERIALS AND METHODS

### Materials

Biochemicals were purchased from Fisher Scientific (Atlanta, GA, USA), Bio-Rad (Hercules, CA, USA), and Sigma-Aldrich (St. Louis, MO, USA). Oligonucleotides and DNA Sanger sequencing services were purchased from Eurofins Genomics (Louisville, KY, USA). Phusion High-Fidelity DNA Polymerase, XhoI, BamHI, DpnI, and 2x Quick Ligase were purchased New England Biolabs (NEB) (Ipswich, MA, USA). DNA fragments were isolated using NEB Monarch PCR & DNA Cleanup Kit or DNA Gel Extraction Kit (Ipswich, MA, USA).

### Plasmid construction and generation of deletion strains

Strains, primers, and plasmids used in this study are listed in **Tables 1-3**. Genomic DNA for plasmid strain construction was extracted from *H. volcanii* by a DNA spooling method (22). The pTA131-based pre-deletion plasmids were generated by restriction enzyme digestion and ligation of a DNA fragment containing the flanking regions 500 bp 5’ (upstream) and 3’ (downstream) of the gene of interest. The knock-out plasmids were constructed by inverse PCR and gel extracted, then treated with DpnI and PCR clean up before adding KLD enzyme mix according to manufacturer’s instruction (New England Biolabs, Ipswich, MA). Plasmids were transformed into *Escherichia coli* Top10, *E. coli* GM2163, and *H. volcanii* H1207. Deletion plasmids to generate *Δelp3* mutations were transformed into *H. volcanii* H1207, KT08, KT09, to generate KT10, KT17, and KT18, respectively.

**Table 1.** List of plasmids used for *H. volcanii* gene deletions in this study.

| Plasmid | Description | Source |
| --- | --- | --- |
| pTA131 | Amp<sup>r</sup>; pBluescript II containing P<sub>fdx</sub>-pyrE2 | (23) |
| pJAM4009 | Amp<sup>r</sup>; pTA131 carrying *pat1* and ~500 bp flanking sequence (pre-KO plasmid) | This study |
| pJAM4013 | Amp<sup>r</sup>; pJAM4009 with $\Delta pat1$ (deletion plasmid) | This study |
| pJAM4010 | Amp<sup>r</sup>; pTA131 carrying *pat2* and ~500 bp flanking sequence (pre-KO plasmid) | This study |
| pJAM4014 | Amp<sup>r</sup>; pJAM4010 with $\Delta pat2$ (deletion plasmid) | This study |
| pJAM4463 | Amp<sup>r</sup>; pTA131 carrying *elp3* and ~500 bp flanking sequence (pre-KO plasmid) | This study |
| pJAM4464 | Amp<sup>r</sup>; pJAM4463 with $\Delta elp3$ (deletion plasmid) | This study |

**Table 2.** List of strains generated and used in this study.

| Strain | Description | Source |
| --- | --- | --- |
| DS70 | wild-type isolate DS2 cured of plasmid pHV2 | (32) |
| H26 | DS70 $\Delta pyrE2$ | (23) |
| H1207 | H26 $\Delta pyrE2$ *pitA*<sub>Nph</sub> $\Delta mrr$ | (33) |
| KT08 | H1207 $\Delta pat1$ | This study |
| KT09 | H1207 $\Delta pat2$ | This study |
| KT10 | H1207 $\Delta elp3$ | This study |
| KT11 | H1207 $\Delta pat1$ $\Delta pat2$ | This study |
| KT17 | H1207 $\Delta pat1$ $\Delta elp3$ | This study |
| KT18 | H1207 $\Delta pat2$ $\Delta elp3$ | This study |

**Table 3.**
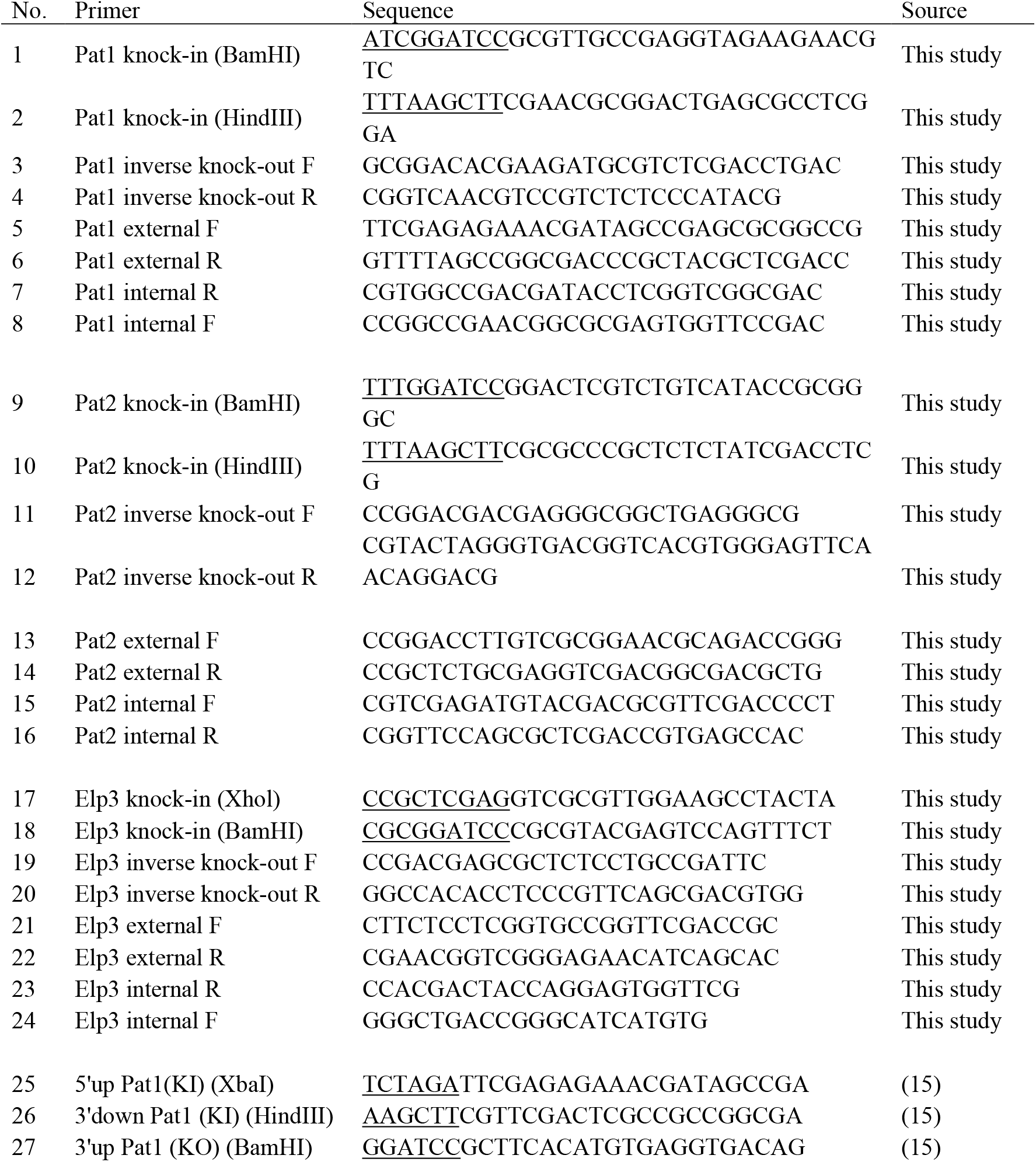
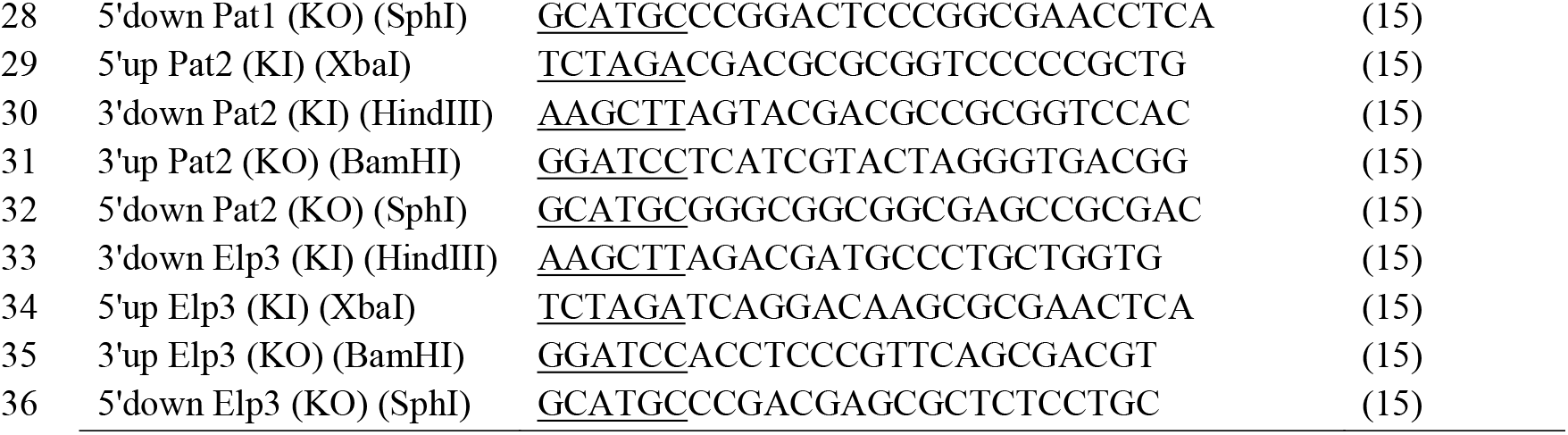
List of primers used in this study compared to Altman-Price and Mevarech (15).

The *H. volcanii* deletion mutants were generated by the *pyrE2* pop-in pop-out homologous recombination method as described (26) with the following modifications. Transformants were plated on Hv-Ca^+^ during the pop in stage, and successful plasmid integration was determined via PCR screening. Positive colonies were grown in 5 mL ATCC974 supplemented with 50 μg/mL 5-FOA (diluted throughout this study from a 50 mg/mL 5-FOA stock dissolved in DMSO) in 13 × 100 mm culture tubes and grown in the dark at 42 °C with orbital shaking at 200 rpm for 3-4 days. Dilution series (10^−3^ to 10^−6^) of this culture was plated on ATCC974 supplemented with 50 μg/mL 5-FOA and 1.5 % (w/v) agar. Genome deletion was determined by PCR screening. Colonies displaying deletion were streaked for isolation four additional times on ATCC974 supplemented with 50 μg/mL 5-FOA. This technique was employed to generate single and double deletions of *pat1* (HVO_1756), *pat2* (HVO_1821), and *elp3* (HVO_2888). Final deletion was monitored by PCR with check primers containing 5’ and 3’ flanking regions 700 bp 5’ (upstream) and 3’ (downstream) of the gene of interest, as well as 300-400 bp internally.

### DNA extraction for genome sequencing

Strains were inoculated from 20 % (v/v) glycerol stocks (−80 °C) onto ATCC974 medium supplemented with 1.5 % (w/v) agar and grown at 42 °C for 5 days. The glycerol stocks were generated by performing a 1:4 dilution of stationary phase cultures with a solution of 80 mL 100 % glycerol supplemented with 20 mL of 30 % salt H_2_O and 0.2 mL of 0.5 M CaCl_2_. From the plates, a single colony was cultured with 5 mL ATCC974 medium in 13 × 100 mm culture tubes and grown until OD_600_ 0.6. To a 2 mL Eppendorf tube, 2 mL of cell culture was pelleted and frozen at 80 °C until further use. Genomic DNA was extracted by ThermoScientific GeneJET Genomic DNA Purification Kit (Atlanta, GA, USA) according to manufacture’s instructions and eluted in H_2_O. Illumina Whole Genome Sequencing (200 Mbp) was performed with variant calling using the reference *H. volcanii* DS2 (NCBI accession NC_013967.1), an Illumina DNA Prep Kit, and the tagmentation method for library preparation (SeqCenter, Pittsburgh, PA).

### PCR Deletion Screening

PCR was performed with Phusion High-Fidelity DNA polymerase according to manufactures instructions with 10 sec extension phases for 25× cycles for all internal primers. The external primers were performed with 1.5 min for *elp3* and 30 sec for *pat1* and *pat2* extension phase for 25× cycles (T100 ThermoCycler, Bio-Rad). PCR products were compared to GeneRuler 1 kb plus DNA ladder (cat# SM1331, ThermoFisher, Waltham, MA, USA). DNA fragments were separated by 0.8 % (w/v) agarose gel supplemented with 0.0025 % (v/v) ethidium bromide electrophoresis (90 V and 30 min) in 1× TAE buffer (0.001 % (v/v) ethidium bromide, 40 mM Tris, 20 mM acetic acid, 1mM EDTA, pH 8.0). DNA agarose gels were imaged using the iBright FL 1000 imaging system (ThermoFisher) on the nucleotide setting.

### Growth curve analysis

Strains were inoculated from 20 % (v/v) glycerol stocks (−80 °C) onto ATCC974 rich medium supplemented with 1.5 % (w/v) agar. Plates were incubated at 42 °C for 5 days. Single colonies were cultured with 5 mL ATCC974 medium in rotating culture tubes (13 × 100 mm). Cells were grown to log phase (OD_600_ 0.6-0.8) and sub-cultured to OD_600_ 0.02 with 5 mL ATCC974 medium and grown to log phase. Cells were sub-cultured again to OD_600_ 0.02 with 1 mL ATCC974 medium in 1.5 mL Eppendorf tubes and briefly (5-10 min) incubated at 42 °C prior to aliquoting. In a 96-well CellPro cell culture plate (Alkali Scientific, FL), 150 µL of subculture was aliquoted into six replicate wells. Using the BioTek Epoch 2 microplate reader and Gen5 software (Agilent, Santa Clara, CA), cell growth was measured as follows: OD_600_ was measured every 15 min for 99 h, with aeration (double orbital continuous shaking), and temperature setpoint 42 °C. No inoculum controls were included to assess potential background signals from the medium alone.

### Data deposition

Whole genome sequencing data for this project was submitted to the National Center for Biotechnology Information (NCBI) Sequence Read Archive (SRA) and can be found under BioProject ID PRJNA1222392 and BioSample IDs SAMN46780405 and SAMN46780406.

## RESULTS

### Single and double mutations of *pat1, pat2* and *elp3* in *H. volcanii*

The essentiality of the *H. volcanii* histone lysine acetyltransferase (HAT) gene homologs was re-evaluated using a pop-in/pop-out method (23) with the strategy to generate markerless deletions of the genes of interest. The plasmids used to target *pat1* (pJAM4013), *pat2* (pJAM4014), and *elp3* (pJAM4464) for deletion (**Table 1**) were transformed into the *H. volcanii* parent strain H1207 to generate single-gene deletions. Subsequently, the *elp3* deletion plasmid was introduced into the single mutant strains KT08 (H1207 *Δpat1*) and KT09 (H1207 *Δpat2*) to construct double mutants. The final strains generated are summarized in **Table 2**. Selection for the *pyrE2* marker was conducted on Hv-Ca^+^ (uracil minus) minimal medium, followed by counter-selection on ATCC974 rich medium supplemented with 5-fluoroorotic acid (5-FOA). Each strain underwent counterselection through four consecutive isolation streaks with PCR screening at each step to monitor successful deletion of the target gene. To further validate the deletions, the strains were recovered from 20 % (v/v) glycerol stocks stored at -80 °C and plated onto ATCC974 rich medium to confirm the absence of the wild type gene. Successful deletion was monitored by PCR analysis using external and internal gene-specific primers (**Fig. 1**). The PCR results demonstrated generation of both single and double mutants of *Δpat1, Δpat2*, and *Δelp3*, with no PCR-detectable wild type signal present.

**Figure 1:**
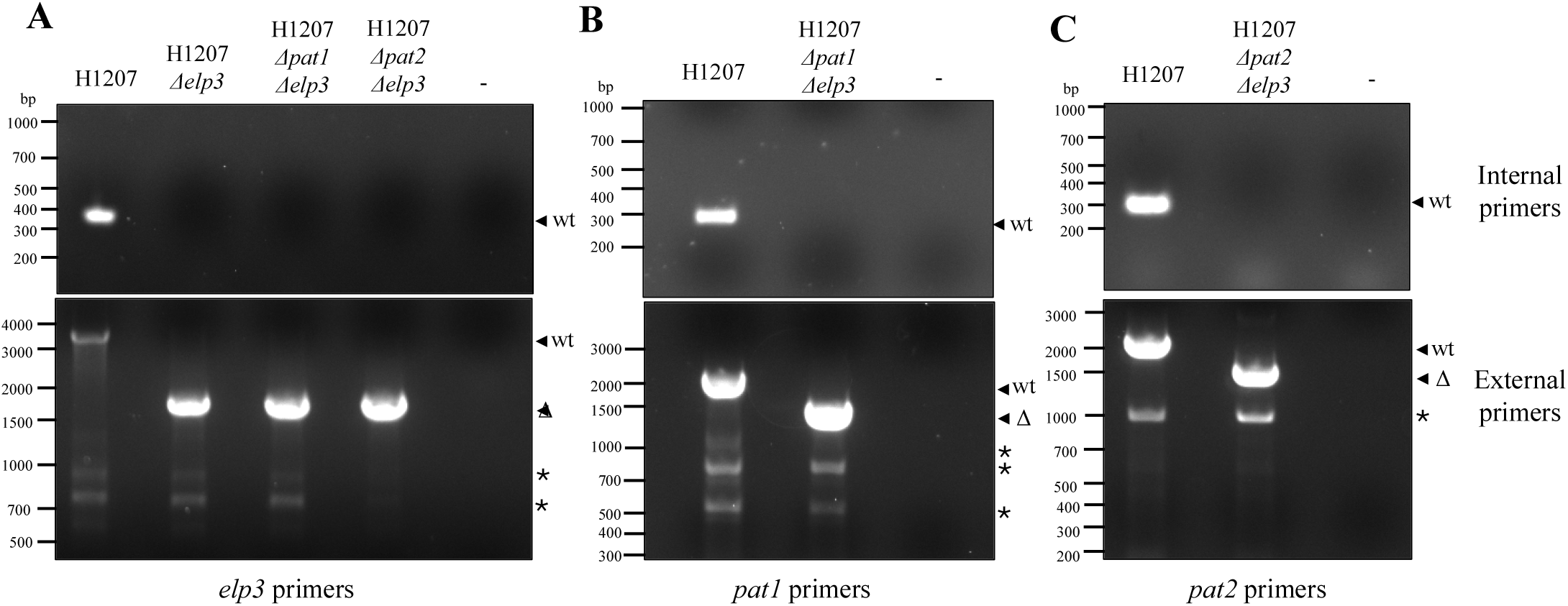
*H. volcanii* strains with single and combined deletions of *elp3* with *pat1* or *pat2* are viable. Parent (H1207) and *Δelp3, Δpat1 Δelp3*, and *Δpat2 Δelp3* mutant strains demonstrated by PCR screen. Cells were inoculated from 20 % (v/v) glycerol stocks (−80 °C) onto ATCC974 medium supplemented with 1.5 % (w/v) agar. A single isolated colony was transferred to a 0.5 mL microcentrifuge tube with 50 µL of sterile H_2_O, heated to 95 °C for 10 min, and cooled at 4 °C for 5 min. The cell material was used for PCR and sterile H_2_O was used for the negative control (−). Deletion of the *elp3* (A), *pat1* (B), and *pat2* (C) genes was monitored by PCR with internal primers (upper panels) and external primes (lower panels), as indicated. The expected sizes for the internal primers were as follows: *elp3* at 400 bp, *pat1* at 302 bp, and *pat2* at 312 bp. The internal primer annealing temperatures were as follows: *elp3* 68 °C, *pat1* 74 °C and *pat2* 74 °C. The expected sizes for the external primers were as follows: *elp3* parent expected 3401 bp vs deletion 1748 bp, *pat1* parent expected 2023 bp vs deletion 1506 bp, *pat2* parent expected 2188 bp vs deletion 1654 bp. The external primer annealing temperatures were as follows: *elp3* 67 °C, *pat1* 74 °C and *pat2* 74 °C. PCR products are labeled on right: *, non-specific; wt, parent; Δ, mutant.

### Whole genome sequencing confirms deletion of *pat2* and *elp3*

To further support the PCR-based screening and verify the successful deletion of the *pat2* and *elp3* genes in a single strain, whole-genome sequencing was performed. Genomic DNA from H1207 and KT18 (H1207 *Δpat2 Δelp3*) was sequenced and analyzed through variant calling against the reference genome (*H. volcanii* DS2; NCBI accession NC_013967.1) (**Supplemental Dataset 1**). The DNA sequence analysis confirmed the successful deletion of 534 bp of *pat2* (GNAT family N-acetyltransferase HVO_RS13445) and 1653 bp of *elp3* (tRNA (uridine 34) 5-carboxymethylaminomethyl modification radical SAM/GNAT enzyme HVO_RS18670) in the KT18 mutant compared to the H1207 parent. According to NCBI (24), *pat2* is 555 bp and *elp3* is 1,659 bp. Additional mutations unique to KT18, relative to H1207, were identified. These additional mutations were insertions at genomic positions 1,697,788 +C, 1,697,869 +A, and 1,697,871 +TGCTCAG that were associated with the ISH3-like element ISHvo20 family transposase pseudogene (*hvo_RS20195*). These mutations within a pseudogene are unlikely to confer suppressor effects during *Δpat2 Δelp3* mutagenesis and are more likely the result of random genomic DNA drift (25). Sequencing coverage for H1207 and H1207 *Δpat2 Δelp3* was 374× and 404×, respectively. Coverage was calculated as the read length (paired-end 150 bp reads) multiplied by the total number of reads for the sample, divided by the genome size (∼2.8 Mbp).

### Phenotypic analysis reveals *Δelp3* single and double mutants with *Δpat1* or *Δpat2* are viable but have growth defects

While the mutation of *elp3*, either alone or in combination with *Δpat1* or *Δpat2*, was not lethal, it did reduce the growth of these mutants compared to the parent. When OD_600_ was used to monitor growth in ATCC974 medium, all three mutant strains (*Δelp3, Δpat1 Δelp3* and *Δpat2 Δelp3*) displayed reduced growth rates, increased doubling times, and lower Area Under the Curve (AUC) values when compared to the parent (H1207) (**Fig. 2**). Furthermore, the *Δpat2 Δelp3* mutant grew somewhat slower than *Δelp3* alone or in combination with *Δpat1*. The parent H1207 was observed to have a doubling time of 4.27 h and growth rate (*μ*) at 0.162 h^−1^, compared to the *Δpat2 Δelp3* mutant which had longer doubling time of 5.08 h and 16% reduced growth rate at 0.136 h^−1^. The *Δelp3* mutant was found to have similar doubling time (4.84 - 4.85 h) and growth rate (0.143 h^−1^) to the *Δpat1Δelp3* mutant, with both strains displaying a 12 % reduction in growth rate compared to the parent. Overall, these findings suggest that *elp3* and *pat2* influence cellular fitness but are not a synthetic lethal gene pairs.

**Figure 2:**
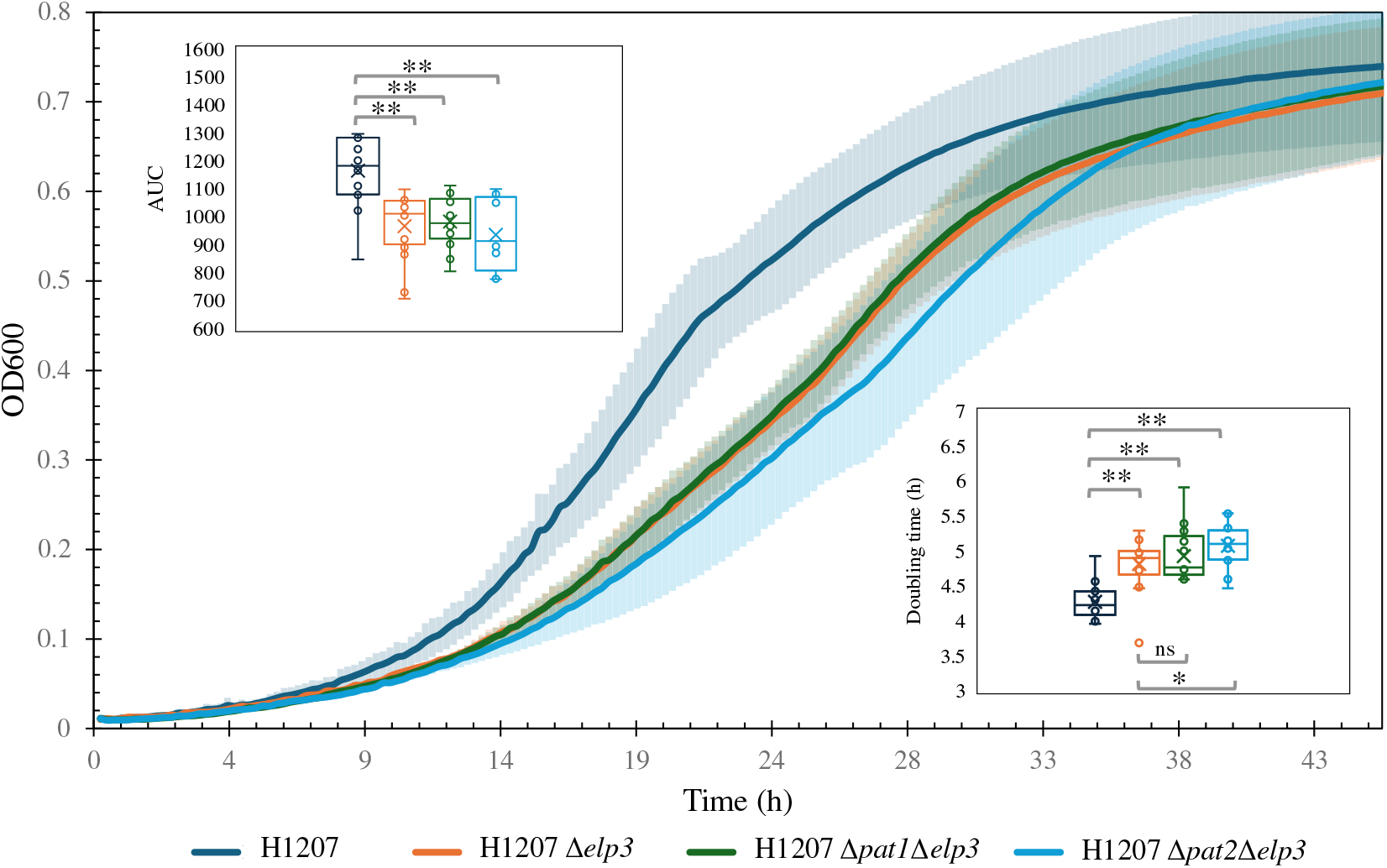
Strains with single and combined deletions of *elp3* with *pat1* or *pat2* are viable but exhibit reduced growth compared to the parent strain (H1207). *H. volcanii* strains were inoculated from 20 % (v/v) glycerol stocks (−80 °C) onto ATCC974 rich medium plates and incubated for 5 days at 42 °C. Isolated colonies were transferred to 5 mL of ATCC 974 and grown to log phase at 42 °C in rotating culture tubes (13×100 mm). Cells were sub-cultured into 5 mL of fresh media at OD_600_ 0.02 and incubated until log phase at 42 °C. For growth monitoring, cells were further sub-cultured into microtiter plates (96-well, OD_600_ of 0.02) and growth was measured at OD_600_ at 15-min intervals using an Epoch 2 Biotech microtiter plate reader at 42 °C with aeration (double orbital continuous shaking). The growth curve represents 3 technical replicates and 6 biological replicates each. No-inoculum control was used as a blank. The experiment was reproducible. ATCC974 medium, pH 6.8, was composed per liter of 125 g NaCl, 50 g MgCl_2_ ·6H_2_O, 5 g K_2_SO_4_, 0.134 g CaCl_2_ ·2H_2_O, 5 g tryptone, and 5 g yeast extract. A student’s t-test was used to determine the statistical significance (p-value <0.005, **) of the area under the curve (AUC) average values calculated over the 46.5 h time course for the parent (H1207, AUC 1170) compared to *Δelp3* (p-value 5.54 × 10^−6^, AUC 983), *Δpat1Δelp3* (p-value 6.20 × 10^−6^, AUC 983), and *Δpat2Δelp3* (p-value 2.25 × 10^−6^, AUC 936) mutant strains.

## DISCUSSION

A previous study (15) aimed to generate single and double mutants of the three lysine acetyltransferase homologs *pat1, pat2*, and *elp3* of *H. volcanii*. With that study (15), the *Δpat2 Δelp3* mutant was not successfully generated, deeming that the deletion of both genes was considered a synthetic lethal due to the potential of their products sharing mutual targets. In this study, we show that *H. volcanii* strains with *elp3* mutations combined with *pat1* or *pat2* deletions remain viable. Whole-genome sequencing further validates the successful double deletion of *pat2* and *elp3* at the genomic level and provides a comprehensive profile of the engineered strains, supporting their suitability for further exploration of lysine acetyltransferase function.

This study also highlights differences between the deletions observed in the current study and those reported in the previous study (15) (**Fig. 3)**. To facilitate deletions, the previous study integrated selection markers on the genome to generate the *Δpat2::hdrB* and *Δelp3::leuB* mutations. The *hdrB* and *leuB* cassettes were placed between the *pat2* and *elp3* gene deletion regions, respectively. This strategy relies on genomic recombination events with the markers adding more selective pressure, allowing for mutations to be more readily isolated (23).

**Figure 3:**
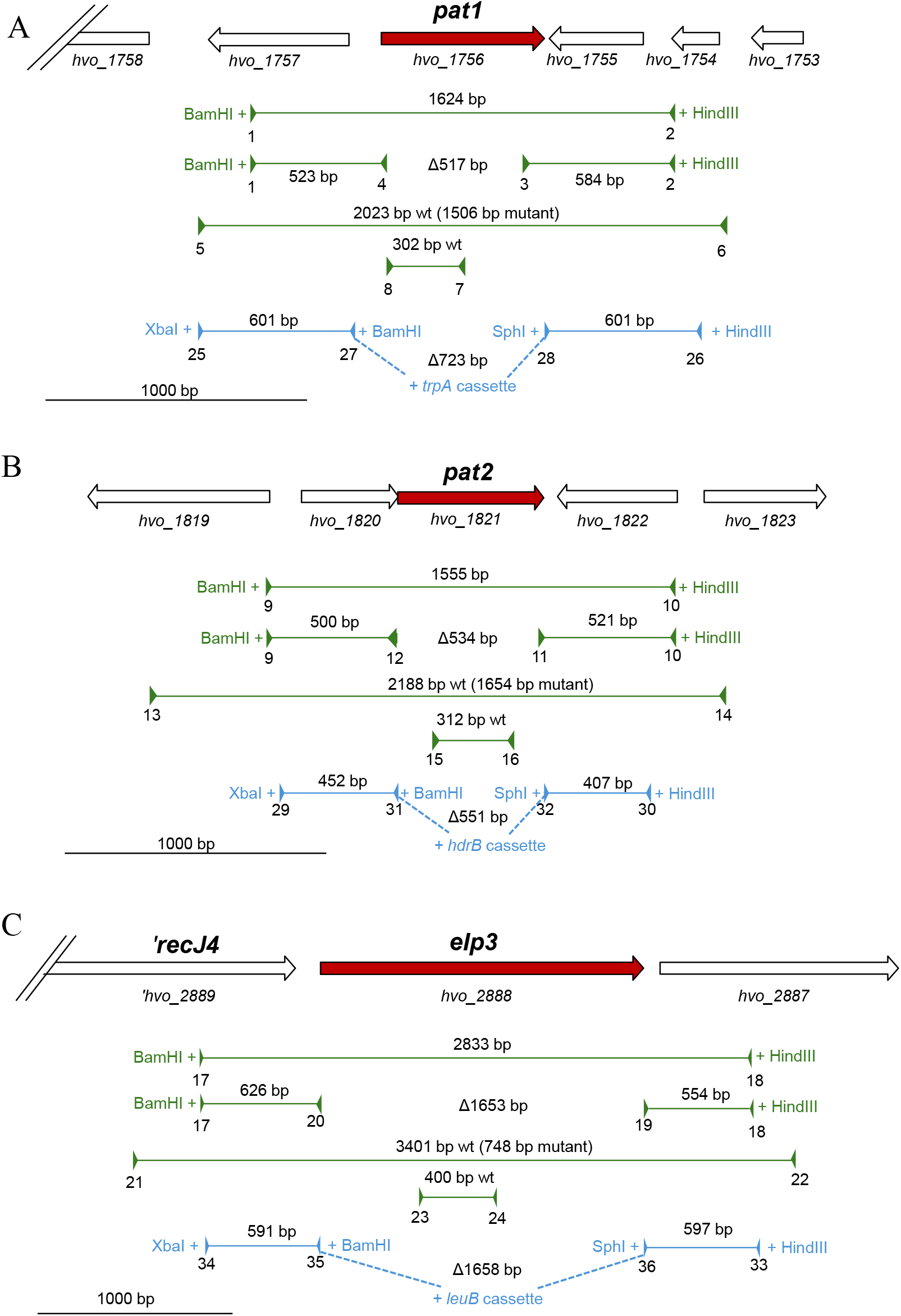
Strategies used to delete the *pat1, pat2*, and *elp3* genes in this study (green) compared to the past work by Altman-Price and Mevarech (15) (blue). The gene of interest is highlighted in red, and neighboring genes are highlighted in white. Plasmids and primers used for mutant strain construction and screening are listed in **Tables 2-3**, with corresponding primer numbers indicated. The pop-in/pop-out method (23) was employed in both studies with the following modifications as outlined below. In this study, the pre-deletion plasmids were generated by inserting the gene of interest with 5’ and 3’ flanking regions into the BamHI and HindIII sites of vector pTA131. The resulting pre-deletion plasmids were used as DNA template to generate the deletion plasmid by inverse PCR. Screening for the mutant strains was performed by PCR with external and internal primers. By contrast, Altman-Price and Mevarech generated the pre-deletion plasmids by inserting the 5’ and 3’ flanking regions of the gene of interest into the XbaI and HindIII sites of pTA131 and including BamHI and SphI sites in this process. The final mutagenesis plasmid was generated by inserting the selection markers *leuB, hdrB*, and *trpA* into the BamHI and SphI sites. The following clarifies the mutagenesis strategy for each gene targeted with primers indicated in parenthesis: For *pat1* (A), a pre-deletion plasmid was generated by inserting a 1,624 bp *pat1* region (1 & 2) into the BamHI and HindIII sites of vector pTA131. The corresponding deletion plasmid was generated by inverse PCR (3 & 4), resulting in a 517 bp deletion (*Δpat1*_*24-539*_). Screening for the deletion was performed using external primers (5 & 6) and internal primers (7 & 8). Altman-Price and Mevarech (15) generated the *pat1* pre-deletion plasmid by inserting two PCR products (25 & 26, 27 & 28) into the XbaI and HindIII sites of pTA131. The *trpA* cassette was inserted into the BamHI and SphI sites of this plasmid to generate the deletion plasmid that would result in a 723 bp deletion with a *trpA* insertion (*Δintergenic*_*-99 to -1*_ *Δpat1*_*1-624*_::*trpA*). GenBank: *pat1* corresponds to CP001956.1: 1625273-1625896 complement; this study generated a deletion of CP001956.1: 1625357-1625873; Altman-Price and Mevarech generated a deletion of CP001956.1: 1625273-1625896. For *pat2* (B), a pre-deletion plasmid was generated by inserting a 1,555 bp *pat2* region (9 & 10) into the BamHI and HindIII sites of vector pTA131. The corresponding deletion plasmid was generated by inverse PCR (11 & 12), resulting in a 534 bp deletion (*Δhvo_1820*_*366-369*_ *Δ pat2*_*1-534*_). Screening for the deletion was performed using external primers (13 & 14) and internal primers (15 & 16). Altman-Price and Mevarech generated the *pat2* pre-deletion plasmid by inserting two PCR products (29 & 30, 31 & 32) into the XbaI and HindIII sites of pTA131. The *hdrB* cassette was inserted into the BamHI and SphI sites of this plasmid to generate the deletion plasmid that would result in a 551 bp deletion and *hdrB* insertion (*Δpat2*_*4-555*_::*hdrB*). GenBank: *pat2* corresponds to CP001956.1: 1683261-1683815; this study generated a deletion of CP001956.1: 1683261-1683794; Altman-Price and Mevarech generated a deletion of CP001956.1: 1625273-1625896. For *elp3* (C), the pre-deletion plasmid was generated by inserting a 2,833 bp *elp3* region (17 & 18) into the BamHI and HindIII sites of vector pTA131. The corresponding deletion plasmid was generated by inverse PCR (19 & 20), resulting in a 1653 bp deletion (*Δelp3*_*6-1659*_). Screening for the deletion was performed using external primers (21 & 22) and internal primers (23 & 24). Altman-Price and Mevarech generated the *elp3* pre-deletion plasmid by inserting two PCR products (33 & 34, 35 & 36) into the XbaI and HindIII sites of pTA131. The *leuB* cassette was inserted into BamHI and SphI sites of this plasmid to generate the deletion plasmid that would result in a 1,658 bp deletion and *leuB* insertion (*Δelp3*_*2-1659*_::*leuB*). GenBank: *elp3* corresponds to CP001956.1: 2726318-2727976 complement; this study generated a deletion of CP001956.1: 2726318-2727970; Altman-Price and Mevarech generated a deletion of CP001956.1: 2726318-2727970.

However, a disadvantage of this approach is the insertion of selection markers into the genome, which can potentially cause distal effects if placed in the middle of an operon or near small open reading frames near neighboring genes. The inverse *pat2* deletion plasmid used in the current study includes a 4-nucleotide deletion at the 5’ end of *hvo_1820*, encoding a universal stress protein A domain (UspA), thus generating the mutant *Δhvo_1820*_*366-369*_ *Δpat2*_*1-534*_. Additionally, the inverse deletion strategy for *pat2* leaves the last 6 amino acids of the C-terminus and the TGA stop codon intact (**Fig. 3**). The mutation of *hvo_1820* may have facilitated the deletion of *pat2* from KT10 by altering stress response pathways and regulation of *pat2*, potentially reducing the resistance to genomic modifications.

We were unsuccessful in generating *Δelp3 Δpat1 Δpat2* triple mutants. The introduction of the *elp3* (pJAM4464) deletion plasmid into KT11 (H1207 *Δpat1 Δpat2*) was achieved up to the final stages of counterselection, as the last 3–4 streaks resulted in unsuccessful isolation between wild type and targeted deletion. Similarly, attempts to introduce the smaller *Δpat1* plasmid (pJAM4013) into KT18 (H1207 *Δpat2 Δelp3*) and the *Δpat2* plasmid (pJAM4014) into KT17 (H1207 *Δpat1 Δelp3*) to generate a triple mutant faced challenges, with wild type being the predominant allele during PCR screening. The polyploid nature of *H. volcanii* (26) is suggested to be a potential factor with generating the triple mutant, resulting in incomplete genome deletion. Polyploidy, presences of multiple copies of the genome, facilitates homologous recombination by providing wild-type templates for the repair of double-strand breaks (27-29).

Polyploidy may also result in wild type trace copies in the cell, if not all genome copies have been successfully mutated. Triple mutations could alternatively be attempted to be generated by introducing the gene of interest on a plasmid or under the control of a tryptophan inducible promoter via conditional depletion, as seen successful for other *H. volcanii* genes (30, 31).

While viable, the strains with the *Δelp3* mutation alone or in combination with either *Δpat1* or *Δpat2* were found to exhibit growth impairments compared to the parent. Our results indicate that *elp3* plays an important role in cellular fitness, *pat2* also influences growth, whereas *pat1* has minimal, if any, impact under the conditions tested. The previous study (15) reported that the *Δpat1, Δpat2, Δelp3, Δpat1 Δpat2*, and *Δpat1 Δelp3* mutant strains were not impacted in growth when compared to the H133 parent strain. In that previous study, *H. volcanii* was cultured at the same temperature (42 °C) but used HY rich medium (per liter: 150 g of NaCl, 36.9 g of MgSO_4_·7H_2_O, 5 mM KCl, 1.07 μM MnCl_2_, 5 g yeast extract, and 50 mM Tris-HCl [pH 7.2]). In contrast, ATCC974 medium [pH 6.8] (per liter: 125 g NaCl, 50 g MgCl_2_ ·6H_2_O, 5 g K_2_SO_4_, 0.134 g CaCl_2_ ·2H_2_O, 5 g tryptone, 5 g yeast extract) was used to determine growth rates in this study. HY rich medium contains 2.57 M NaCl compared to ATCC974 at 2.14 M NaCl, a 1.2-fold difference of NaCl concentration. HY medium also includes MnCl2 as a trace element and Tris-HCl for pH buffering, while ATCC974 contains a calcium source (CaCl2) and tryptone, an additional source of peptides and amino acids. ATCC974 medium contains a lower salt concentration, which offers advantages for autoclaving, and may account for the differences including our ability to detect a phenotype and generate the *Δpat2 Δelp3* mutant strain.

In conclusion, we provide strong evidence that *elp3* and *pat2* can be deleted in the same *H. volcanii* strain based on whole genome sequencing. Beyond the targeted deletions, minimal differences between the parent and *Δelp3 Δpat2* mutant were observed suggesting that suppressor mutations are not responsible for our ability to generate this double mutant strain. Elp3 and Pat2, thus, may not share as close a functional relationship as implied by earlier study (15). Our finding is significant as Elp3 is thought to function in acetylation in tRNA modification, while Pat2 likely functions in the lysine acetylation of proteins.

## Acknowledgements

Funds awarded to JMF through U.S. Department of Energy, Office of Basic Energy Sciences, Division of Chemical Sciences, Geosciences and Biosciences, Physical Biosciences Program [DOE DE-FG02-05ER15650] by determine global mechanisms of redox regulation in archaea and the National Institutes of Health [NIH R01 GM057498] by providing evolutionary insight in biological systems.

